## Supplementary Figures for "The adenoviral E1B-55k protein present in HEK293 cells mediates abnormal accumulation of key WNT signaling proteins in large cytoplasmic aggregates"

SUPPLEMENTARY INFORMATION

**Figure S1. Localization of key WNT/β-catenin signaling components in the HEK293T cell line.**

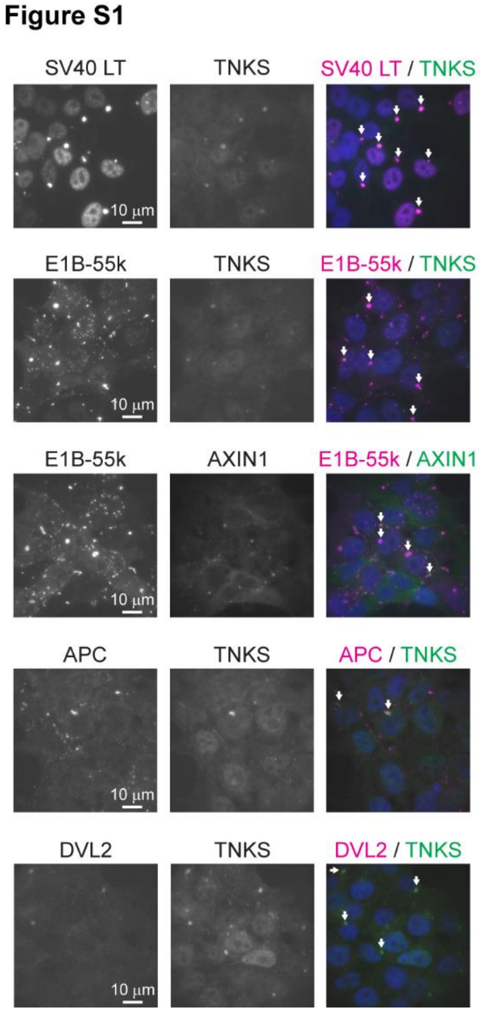


**Figure S1.** Localization of key WNT/β-catenin signaling components in the HEK293T cell line.

IF analysis (WF) of the localization of SV40-Large T antigen ((SV40 LT) not present in HEK293 cells), TNKS, E1B-55k, AXIN1, APC and DVL2 as indicated. In the merged color images co-localization is indicated with arrows and nuclear counterstaining is shown in blue.

**Figure S2. Measurement of E1B-55k protein layer thickness.**

**Figure S2**

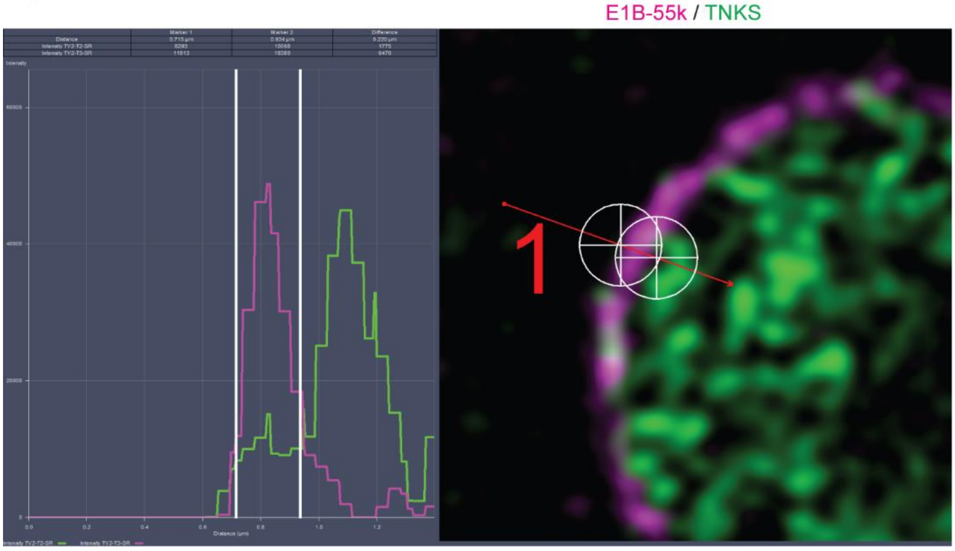


Figure S2. Measurement of E1B-55k protein layer thickness.

The thickness of the E1B-55k protein layer was measured in a SIM image slice using the ZEN black software measurement tool. The histogram to the left shows the intensity plot of the E1B-55k (magenta) and TNKS (green) IF staining over the area covered by the red line in the image to the right. The white vertical lines in the histogram correspond to the center position of the aim crosses in the image. The measured distance between the lines and hence the thickness of the E1B-55k protein layer at the indicated position is 220 nm.

**Figure S3**. **Western blot analysis of E1B-55k protein levels in siRNA treated HEK293 cells.**

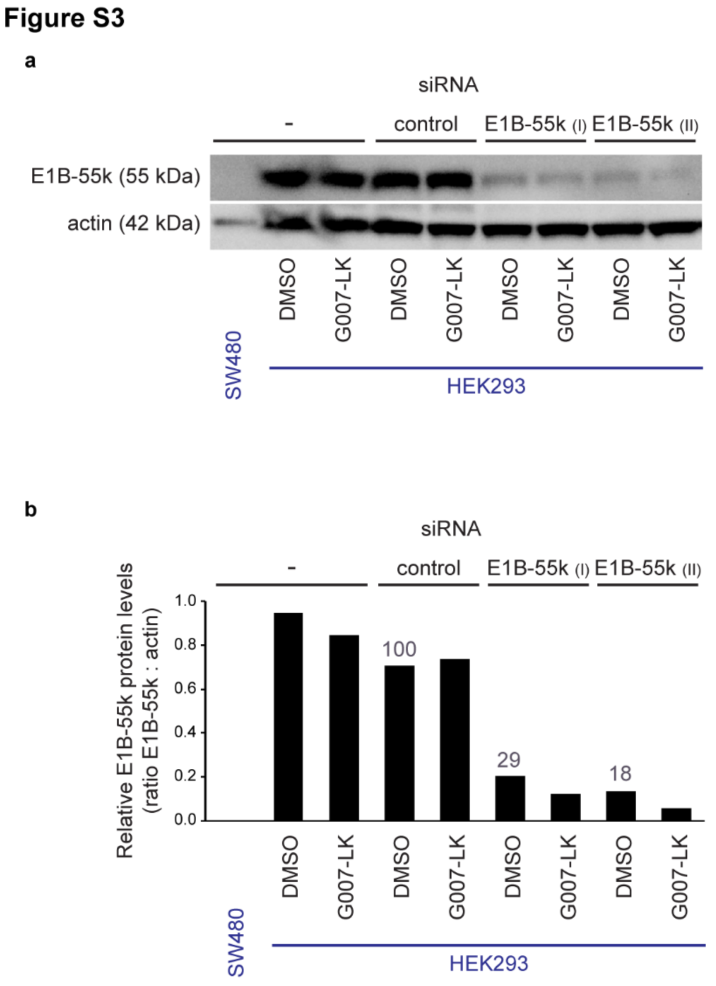


Figure S2. Western blot analysis of E1B-55k protein levels in siRNA treated HEK293 cells.

(a) Western blot analysis of E1B-55k protein levels in untreated (-) and HEK293 cells treated with control siRNA or two independent E1B-55k siRNA constructs (I and II). Both DMSO and G007-LK treated HEK293 cells were analyzed as indicated. As a negative control, protein extracts from a colon carcinoma cell line (SW480), not expressing E1B-55k, was included. Bands for E1B-55k protein and for the actin loading control is shown. (b) Quantification of E1B-55k protein levels as determined by calculating the ratio of the background subtracted intensities of the E1B-55k bands relative to the corresponding actins band in A (Image Lab Software, Bio-Rad). Treatment with E1B-55k siRNA I and II reduced E1B-55k protein levels to 29 % and 18 % respectively, relative to the protein levels in control siRNA treated cells.

**Figure S4. In HEK293 cells with reduced E1B-55k protein levels the distribution of AXIN1, APC and DVL2 is changed from accumulation in aggregates to a more uniform cytoplasmic distribution.**

**Figure S4**

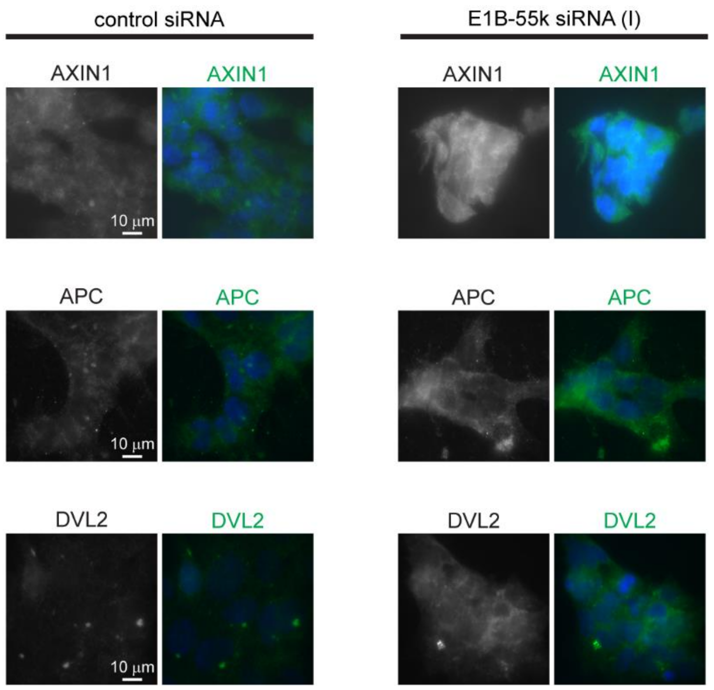


Figure S4. In HEK293 cells with reduced E1B-55k protein levels the distribution of AXIN1, APC and DVL2 is changed from accumulation in aggregates to a more uniform cytoplasmic distribution.

IF analysis (laser WF) of the localization of AXIN1, APC and DVL2 in HEK293 cells treated with control or E1B-55k siRNA. In the merged color images nuclear counterstaining is shown in blue.
