## Supplementary Tables for "The adenoviral E1B-55k protein present in HEK293 cells mediates abnormal accumulation of key WNT signaling proteins in large cytoplasmic aggregates"

**Supplementary Table 1. E1B-55k interacting partners detected in Co-IP experiment 1.**

| Accession | Gene name | Description | Coverage | # Peptides | # PSMs | # Unique Peptides | # Protein Groups | # AAs | MW [kDa] | calc. pI | Beads only control | Mouse IgG control | E1B-55k Ab | Protein FDR Confidence Mascot | Exp. q-value Mascot | Mascot Score | # Peptides Mascot |
|---|---|---|---|---|---|---|---|---|---|---|---|---|---|---|---|---|---|
| Q92878 | RAD50 | DNA repair protein RAD50 OS=Homo sapiens GN=RAD50 PE=1 SV=1 | 27,668 | 34 | 39 | 34 | 1 | 1312 | 153,797 | 6,89 | Not Found | Not Found | High | High | 0,000 | 236,18 | 4 |
| P49959 | MRE11 | Double-strand break repair protein MRE11A OS=Homo sapiens GN=MRE11A PE=1 SV=3 | 35,452 | 21 | 21 | 21 | 1 | 708 | 80,543 | 5,9 | Not Found | Not Found | High | High | 0,000 | 233,41 | 5 |
| Q93008 | USP9X | Probable ubiquitin carboxyl-terminal hydrolase FAF-X OS=Homo sapiens GN=USP9X PE=1 SV=3 | 6,926 | 15 | 16 | 15 | 1 | 2570 | 292,094 | 5,8 | Not Found | Not Found | High | High | 0,000 | 230,37 | 6 |
| P04637 | TP53 | Cellular tumor antigen p53 OS=Homo sapiens GN=TP53 PE=1 SV=4 | 26,209 | 10 | 10 | 10 | 1 | 393 | 43,625 | 6,79 | Not Found | Not Found | High | High | 0,000 | 228,01 | 2 |
| Q96C92 | SDCCAG3 | Serologically defined colon cancer antigen 3 OS=Homo sapiens GN=SDCCAG3 PE=1 SV=3 | 17,931 | 6 | 7 | 6 | 1 | 435 | 47,932 | 5,14 | Not Found | Not Found | High | High | 0,000 | 223,93 | 3 |
| Q8N1N4 | KRT78 | Keratin, type II cytoskeletal 78 OS=Homo sapiens GN=KRT78 PE=2 SV=2 | 11,731 | 6 | 8 | 4 | 1 | 520 | 56,83 | 6,02 | Not Found | Not Found | High | High | 0,000 | 221,95 | 6 |
| Q5JSZ5 | PRRC2B | Protein PRRC2B OS=Homo sapiens GN=PRRC2B PE=1 SV=2 | 1,660 | 3 | 3 | 3 | 1 | 2229 | 242,817 | 8,34 | Not Found | Not Found | High | High | 0,000 | 220,30 | 3 |
| Q9NVI7 | ATAD3A | ATPase family AAA domain-containing protein 3A OS=Homo sapiens GN=ATAD3A PE=1 SV=2 | 7,886 | 5 | 6 | 5 | 1 | 634 | 71,325 | 8,98 | Not Found | Not Found | High | High | 0,000 | 218,38 | 1 |
| Q86SQ0 | PHLDB2 | Pleckstrin homology-like domain family B member 2 OS=Homo sapiens GN=PHLDB2 PE=1 SV=2 | 4,469 | 5 | 5 | 5 | 1 | 1253 | 142,07 | 7,43 | Not Found | Not Found | High | High | 0,000 | 215,79 | 3 |
| P05089 | ARG1 | Arginase-1 OS=Homo sapiens GN=ARG1 PE=1 SV=2 | 18,944 | 4 | 4 | 4 | 1 | 322 | 34,713 | 7,21 | Not Found | Not Found | High | High | 0,000 | 211,97 | 5 |
| O60934 | NBN | Nibrin OS=Homo sapiens GN=NBN PE=1 SV=1 | 4,244 | 3 | 3 | 3 | 1 | 754 | 84,906 | 6,9 | Not Found | Not Found | High | High | 0,000 | 206,41 | 5 |
| Q08188 | TGM3 | Protein-glutamine gamma-glutamyltransferase E OS=Homo sapiens GN=TGM3 PE=1 SV=4 | 4,185 | 2 | 2 | 2 | 1 | 693 | 76,584 | 5,86 | Not Found | Not Found | High | High | 0,000 | 201,22 | 5 |
| P31944 | CASP14 | Caspase-14 OS=Homo sapiens GN=CASP14 PE=1 SV=2 | 13,223 | 3 | 3 | 3 | 1 | 242 | 27,662 | 5,58 | Not Found | Not Found | High | High | 0,000 | 200,38 | 4 |
| A2RUB1 | MEIOC | Meiosis-specific coiled-coil domain-containing protein MEIOC OS=Homo sapiens GN=MEIOC PE=2 SV=3 | 2,206 | 2 | 2 | 2 | 1 | 952 | 107,491 | 7,12 | Not Found | Not Found | High | High | 0,000 | 198,96 | 3 |
| Q8NDV7 | TNRC6A | Trinucleotide repeat-containing gene 6A protein OS=Homo sapiens GN=TNRC6A PE=1 SV=2 | 1,478 | 2 | 2 | 2 | 1 | 1962 | 210,169 | 7,01 | Not Found | Not Found | High | High | 0,000 | 197,88 | 3 |
| O95816 | BAG2 | BAG family molecular chaperone regulator 2 OS=Homo sapiens GN=BAG2 PE=1 SV=1 | 13,744 | 3 | 3 | 3 | 1 | 211 | 23,757 | 6,7 | Not Found | Not Found | High | High | 0,000 | 196,78 | 2 |
| Q5VUA4 | ZNF318 | Zinc finger protein 318 OS=Homo sapiens GN=ZNF318 PE=1 SV=2 | 2,106 | 3 | 3 | 3 | 1 | 2279 | 250,958 | 7,2 | Not Found | Not Found | High | High | 0,000 | 191,71 | 3 |
| O43663 | PRC1 | Protein regulator of cytokinesis 1 OS=Homo sapiens GN=PRC1 PE=1 SV=2 | 3,226 | 2 | 2 | 2 | 1 | 620 | 71,562 | 6,68 | Not Found | Not Found | High | High | 0,000 | 191,62 | 1 |
| Q9UPQ9 | TNRC6B | Trinucleotide repeat-containing gene 6B protein OS=Homo sapiens GN=TNRC6B PE=1 SV=4 | 1,200 | 2 | 2 | 2 | 1 | 1833 | 193,883 | 6,76 | Not Found | Not Found | High | High | 0,000 | 181,99 | 2 |
| Q6Q0C0 | TRAF7 | E3 ubiquitin-protein ligase TRAF7 OS=Homo sapiens GN=TRAF7 PE=1 SV=1 | 1,493 | 1 | 1 | 1 | 1 | 670 | 74,561 | 7,15 | Not Found | Not Found | High | High | 0,000 | 181,43 | 3 |
| Q13751 | LAMB3 | Laminin subunit beta-3 OS=Homo sapiens GN=LAMB3 PE=1 SV=1 | 1,280 | 1 | 1 | 1 | 1 | 1172 | 129,489 | 7,21 | Not Found | Not Found | High | High | 0,000 | 177,27 | 2 |
| Q8WWI1 | LMO7 | LIM domain only protein 7 OS=Homo sapiens GN=LMO7 PE=1 SV=3 | 0,891 | 1 | 1 | 1 | 1 | 1683 | 192,576 | 8,09 | Not Found | Not Found | High | High | 0,000 | 175,38 | 5 |
| Q14999 | CUL7 | Cullin-7 OS=Homo sapiens GN=CUL7 PE=1 SV=2 | 1,767 | 3 | 3 | 2 | 1 | 1698 | 191,04 | 5,87 | Not Found | Not Found | High | High | 0,000 | 174,62 | 4 |
| P25311 | AZGP1 | Zinc-alpha-2-glycoprotein OS=Homo sapiens GN=AZGP1 PE=1 SV=2 | 3,356 | 1 | 1 | 1 | 1 | 298 | 34,237 | 6,05 | Not Found | Not Found | High | High | 0,000 | 169,56 | 3 |
| P84090 | ERH | Enhancer of rudimentary homolog OS=Homo sapiens GN=ERH PE=1 SV=1 | 21,154 | 2 | 2 | 2 | 1 | 104 | 12,251 | 5,92 | Not Found | Not Found | High | High | 0,000 | 168,64 | 4 |
| Q5T749 | KPRP | Keratinocyte proline-rich protein OS=Homo sapiens GN=KPRP PE=1 SV=1 | 7,772 | 3 | 3 | 3 | 1 | 579 | 64,093 | 8,27 | Not Found | Not Found | High | High | 0,000 | 168,29 | 3 |
| O43795 | MYO1B | Unconventional myosin-Ib OS=Homo sapiens GN=MYO1B PE=1 SV=3 | 1,849 | 1 | 1 | 1 | 1 | 1136 | 131,902 | 9,38 | Not Found | Not Found | High | High | 0,000 | 168,12 | 6 |
| O43823 | AKAP8 | A-kinase anchor protein 8 OS=Homo sapiens GN=AKAP8 PE=1 SV=1 | 1,879 | 1 | 1 | 1 | 1 | 692 | 76,061 | 5,15 | Not Found | Not Found | High | High | 0,000 | 167,47 | 5 |
| P06733 | ENO1 | Alpha-enolase OS=Homo sapiens GN=ENO1 PE=1 SV=2 | 3,687 | 2 | 2 | 2 | 1 | 434 | 47,139 | 7,39 | Not Found | Not Found | High | High | 0,000 | 166,22 | 3 |
| O75223 | GGCT | Gamma-glutamylcyclotransferase OS=Homo sapiens GN=GGCT PE=1 SV=1 | 6,915 | 1 | 1 | 1 | 1 | 188 | 20,994 | 5,14 | Not Found | Not Found | High | High | 0,000 | 161,04 | 2 |
| Q9UPN4 | CEP131 | Centrosomal protein of 131 kDa OS=Homo sapiens GN=CEP131 PE=1 SV=3 | 1,477 | 1 | 1 | 1 | 1 | 1083 | 122,075 | 8,69 | Not Found | Not Found | High | High | 0,000 | 160,86 | 4 |
| Q8WVV4 | POF1B | Protein POF1B OS=Homo sapiens GN=POF1B PE=1 SV=3 | 1,868 | 1 | 1 | 1 | 1 | 589 | 68,022 | 6,32 | Not Found | Not Found | High | High | 0,000 | 160,39 | 3 |
| Q9ULV4 | CORO1C | Coronin-1C OS=Homo sapiens GN=CORO1C PE=1 SV=1 | 1,688 | 1 | 1 | 1 | 1 | 474 | 53,215 | 7,08 | Not Found | Not Found | High | High | 0,000 | 158,90 | 3 |
| P14373 | TRIM27 | Zinc finger protein RFP OS=Homo sapiens GN=TRIM27 PE=1 SV=1 | 2,729 | 1 | 1 | 1 | 1 | 513 | 58,452 | 6,21 | Not Found | Not Found | High | High | 0,000 | 157,82 | 3 |
| P52597 | HNRNPF | Heterogeneous nuclear ribonucleoprotein F OS=Homo sapiens GN=HNRNPF PE=1 SV=3 | 4,096 | 1 | 1 | 1 | 1 | 415 | 45,643 | 5,58 | Not Found | Not Found | High | High | 0,000 | 155,60 | 4 |

| | | | | | | | | | | | | | | | | | |
|---|---|---|---|---|---|---|---|---|---|---|---|---|---|---|---|---|---|
| Q9NZT1 | CALML5 | Calmodulin-like protein 5 OS=Homo sapiens GN=CALML5 PE=1 SV=2 | 8,904 | 1 | 1 | 1 | 1 | 146 | 15,883 | 4,44 | Not Found | Not Found | High | High | 0,000 | 153,43 | 6 |
| P48668 | KRT6C | Keratin, type II cytoskeletal 6C OS=Homo sapiens GN=KRT6C PE=1 SV=3 | 52,305 | 35 | 108 | 2 | 1 | 564 | 59,988 | 8 | Not Found | Not Found | Medium | High | 0,001 | 112,04 | 2 |
| Q14574 | DSC3 | Desmocollin-3 OS=Homo sapiens GN=DSC3 PE=1 SV=3 | 2,567 | 2 | 2 | 2 | 1 | 896 | 99,906 | 6,1 | Not Found | Not Found | Medium | High | 0,001 | 108,73 | 1 |
| O75420 | GIGYF1 | PERQ amino acid-rich with GYF domain-containing protein 1 OS=Homo sapiens GN=GIGYF1 PE=1 SV=2 | 2,802 | 2 | 2 | 2 | 1 | 1035 | 114,531 | 5,39 | Not Found | Not Found | Medium | High | 0,001 | 108,60 | 2 |
| Q8IWT3 | CUL9 | Cullin-9 OS=Homo sapiens GN=CUL9 PE=1 SV=2 | 0,874 | 2 | 2 | 1 | 1 | 2517 | 281,049 | 5,45 | Not Found | Not Found | Medium | High | 0,001 | 108,53 | 2 |
| O00159 | MYO1C | Unconventional myosin-Ic OS=Homo sapiens GN=MYO1C PE=1 SV=4 | 1,693 | 2 | 2 | 2 | 1 | 1063 | 121,606 | 9,41 | Not Found | Not Found | Medium | High | 0,001 | 107,00 | 2 |
| P35580 | MYH10 | Myosin-10 OS=Homo sapiens GN=MYH10 PE=1 SV=3 | 0,810 | 1 | 1 | 1 | 1 | 1976 | 228,858 | 5,54 | Not Found | Not Found | Medium | High | 0,001 | 106,08 | 2 |
| O75342 | ALOX12B | Arachidonate 12-lipoxygenase, 12R-type OS=Homo sapiens GN=ALOX12B PE=1 SV=1 | 1,712 | 1 | 1 | 1 | 1 | 701 | 80,304 | 7,64 | Not Found | Not Found | Medium | High | 0,001 | 104,66 | 1 |
| P40939 | HADHA | Trifunctional enzyme subunit alpha, mitochondrial OS=Homo sapiens GN=HADHA PE=1 SV=2 | 2,752 | 1 | 1 | 1 | 1 | 763 | 82,947 | 9,04 | Not Found | Not Found | Medium | High | 0,001 | 104,42 | 2 |
| Q96P63 | SERPINB12 | Serpin B12 OS=Homo sapiens GN=SERPINB12 PE=1 SV=1 | 3,457 | 1 | 1 | 1 | 1 | 405 | 46,247 | 5,53 | Not Found | Not Found | Medium | High | 0,002 | 101,71 | 1 |
| Q9H7T9 | AUNIP | Aurora kinase A and ninein-interacting protein OS=Homo sapiens GN=AUNIP PE=1 SV=1 | 3,361 | 1 | 1 | 1 | 1 | 357 | 40,228 | 7,53 | Not Found | Not Found | Medium | High | 0,002 | 101,47 | 3 |
| Q9UHB6 | LIMA1 | LIM domain and actin-binding protein 1 OS=Homo sapiens GN=LIMA1 PE=1 SV=1 | 1,449 | 1 | 1 | 1 | 1 | 759 | 85,173 | 6,84 | Not Found | Not Found | Medium | High | 0,002 | 95,88 | 2 |
| Q7Z5K2 | WAPL | Wings apart-like protein homolog OS=Homo sapiens GN=WAPL PE=1 SV=1 | 1,092 | 1 | 1 | 1 | 1 | 1190 | 132,863 | 5,44 | Not Found | Not Found | Medium | High | 0,002 | 90,14 | 1 |
| P20930 | FLG | Filaggrin OS=Homo sapiens GN=FLG PE=1 SV=3 | 0,837 | 2 | 2 | 2 | 1 | 4061 | 434,922 | 9,25 | Not Found | Not Found | Medium | High | 0,002 | 90,10 | 2 |
| Q96QB1 | DLC1 | Rho GTPase-activating protein 7 OS=Homo sapiens GN=DLC1 PE=1 SV=4 | 0,654 | 1 | 1 | 1 | 1 | 1528 | 170,485 | 6,4 | Not Found | Not Found | Medium | High | 0,003 | 84,96 | 1 |
| Q15058 | KIF14 | Kinesin-like protein KIF14 OS=Homo sapiens GN=KIF14 PE=1 SV=1 | 1,214 | 2 | 2 | 2 | 1 | 1648 | 186,375 | 7,91 | Not Found | Not Found | Medium | High | 0,003 | 82,61 | 3 |
| Q92841 | DDX17 | Probable ATP-dependent RNA helicase DDX17 OS=Homo sapiens GN=DDX17 PE=1 SV=2 | 1,646 | 1 | 1 | 1 | 1 | 729 | 80,222 | 8,27 | Not Found | Not Found | Medium | High | 0,003 | 79,76 | 2 |
| O60573 | EIF4E2 | Eukaryotic translation initiation factor 4E type 2 OS=Homo sapiens GN=EIF4E2 PE=1 SV=1 | 4,082 | 1 | 1 | 1 | 1 | 245 | 28,344 | 8,88 | Not Found | Not Found | Medium | High | 0,004 | 77,10 | 2 |
| Q8NB66 | UNC13C | Protein unc-13 homolog C OS=Homo sapiens GN=UNC13C PE=2 SV=3 | 0,361 | 1 | 1 | 1 | 1 | 2214 | 250,754 | 5,92 | Not Found | Not Found | Medium | High | 0,004 | 75,84 | 2 |
| O14654 | IRS4 | Insulin receptor substrate 4 OS=Homo sapiens GN=IRS4 PE=1 SV=1 | 1,273 | 1 | 1 | 1 | 1 | 1257 | 133,685 | 8,44 | Not Found | Not Found | Medium | High | 0,004 | 75,77 | 1 |
| Q9NYF8 | BCLAF1 | Bcl-2-associated transcription factor 1 OS=Homo sapiens GN=BCLAF1 PE=1 SV=2 | 1,522 | 1 | 1 | 1 | 1 | 920 | 106,059 | 9,98 | Not Found | Not Found | Medium | High | 0,004 | 74,43 | 2 |
| P01861 | IGHG4 | Ig gamma-4 chain C region OS=Homo sapiens GN=IGHG4 PE=1 SV=1 | 2,752 | 1 | 1 | 1 | 1 | 327 | 35,918 | 7,36 | Not Found | Not Found | Medium | High | 0,004 | 74,19 | 2 |

**Supplementary Table 2. E1B-55k interacting partners detected in Co-IP experiment 2.**

| Accession | Gene name | Description | Coverage | # Peptides | # PSMs | # Unique Peptides | # Protein Groups | # AAs | MW [kDa] | calc. pI | Beads only control | Mouse IgG control | E1B-55k Ab | Protein FDR Confidence Mascot | Exp. q-value Mascot | Mascot Score | # Peptides Mascot |
|---|---|---|---|---|---|---|---|---|---|---|---|---|---|---|---|---|---|
| Q92878 | RAD50 | DNA repair protein RAD50 OS=Homo sapiens GN=RAD50 PE=1 SV=1 | 33,99390244 | 41 | 50 | 41 | 1 | 1312 | 153,797 | 6,89 | Not Found | Not Found | High | High | 0,000 | 1521,70 | 41 |
| Q93008 | USP9X | Probable ubiquitin carboxyl-terminal hydrolase FAF-X OS=Homo sapiens GN=USP9X PE=1 SV=3 | 13,46303502 | 24 | 26 | 24 | 1 | 2570 | 292,094 | 5,8 | Not Found | Not Found | High | High | 0,000 | 711,18 | 24 |
| P49959 | MRE11 | Double-strand break repair protein MRE11 OS=Homo sapiens GN=MRE11 PE=1 SV=3 | 26,83615819 | 14 | 18 | 14 | 1 | 708 | 80,543 | 5,9 | Not Found | Not Found | High | High | 0,000 | 455,45 | 14 |
| P04637 | TP53 | Cellular tumor antigen p53 OS=Homo sapiens GN=TP53 PE=1 SV=4 | 25,95419847 | 7 | 12 | 4 | 1 | 393 | 43,625 | 6,79 | Not Found | Not Found | High | High | 0,000 | 436,61 | 7 |
| Q5JSZ5 | PRRC2B | Protein PRRC2B OS=Homo sapiens GN=PRRC2B PE=1 SV=2 | 1,973979363 | 3 | 4 | 3 | 1 | 2229 | 242,817 | 8,34 | Not Found | Not Found | High | High | 0,000 | 290,18 | 3 |
| Q86SQ0 | PHLDB2 | Pleckstrin homology-like domain family B member 2 OS=Homo sapiens GN=PHLDB2 PE=1 SV=2 | 8,379888268 | 8 | 8 | 8 | 1 | 1253 | 142,07 | 7,43 | Not Found | Not Found | High | High | 0,000 | 216,19 | 8 |
| O60934 | NBN | Nibrin OS=Homo sapiens GN=NBN PE=1 SV=1 | 10,87533156 | 7 | 7 | 7 | 1 | 754 | 84,906 | 6,9 | Not Found | Not Found | High | High | 0,000 | 189,02 | 7 |
| Q9Y5X1 | SNX9 | Sorting nexin-9 OS=Homo sapiens GN=SNX9 PE=1 SV=1 | 16,80672269 | 6 | 6 | 6 | 1 | 595 | 66,55 | 5,58 | Not Found | Not Found | High | High | 0,000 | 188,73 | 6 |
| Q9H6S0 | YTHDC2 | Probable ATP-dependent RNA helicase YTHDC2 OS=Homo sapiens GN=YTHDC2 PE=1 SV=2 | 4,615384615 | 4 | 4 | 2 | 1 | 1430 | 160,147 | 8,4 | Not Found | Not Found | High | High | 0,000 | 143,30 | 4 |
| P40939 | HADHA | Trifunctional enzyme subunit alpha, mitochondrial OS=Homo sapiens GN=HADHA PE=1 SV=2 | 8,125819135 | 4 | 4 | 4 | 1 | 763 | 82,947 | 9,04 | Not Found | Not Found | High | High | 0,000 | 133,24 | 4 |
| P25054 | APC | Adenomatous polyposis coli protein OS=Homo sapiens GN=APC PE=1 SV=2 | 1,793879705 | 4 | 4 | 4 | 1 | 2843 | 311,455 | 7,8 | Not Found | Not Found | High | High | 0,000 | 132,09 | 4 |
| Q7L804 | RAB11FIP2 | Rab11 family-interacting protein 2 OS=Homo sapiens GN=RAB11FIP2 PE=1 SV=1 | 8,0078125 | 3 | 3 | 3 | 1 | 512 | 58,243 | 9,32 | Not Found | Not Found | High | High | 0,001 | 115,78 | 3 |
| Q7Z5K2 | WAPL | Wings apart-like protein homolog OS=Homo sapiens GN=WAPL PE=1 SV=1 | 3,697478992 | 2 | 2 | 2 | 1 | 1190 | 132,863 | 5,44 | Not Found | Not Found | High | High | 0,001 | 114,53 | 2 |
| Q02413 | DSG1 | Desmoglein-1 OS=Homo sapiens GN=DSG1 PE=1 SV=2 | 4,671115348 | 3 | 4 | 3 | 1 | 1049 | 113,676 | 5,03 | Not Found | Not Found | High | High | 0,002 | 106,87 | 3 |
| Q13428 | TCOF1 | Treacle protein OS=Homo sapiens GN=TCOF1 PE=1 SV=3 | 2,419354839 | 3 | 3 | 3 | 1 | 1488 | 152,015 | 9,04 | Not Found | Not Found | High | High | 0,002 | 106,12 | 3 |
| P14373 | TRIM27 | Zinc finger protein RFP OS=Homo sapiens GN=TRIM27 PE=1 SV=1 | 5,068226121 | 2 | 2 | 2 | 1 | 513 | 58,452 | 6,21 | Not Found | Not Found | High | High | 0,002 | 105,73 | 2 |
| Q96C92 | SDCCAG3 | Serologically defined colon cancer antigen 3 OS=Homo sapiens GN=SDCCAG3 PE=1 SV=3 | 6,436781609 | 2 | 2 | 2 | 1 | 435 | 47,932 | 5,14 | Not Found | Not Found | High | High | 0,002 | 97,60 | 2 |
| O95816 | BAG2 | BAG family molecular chaperone regulator 2 OS=Homo sapiens GN=BAG2 PE=1 SV=1 | 17,53554502 | 3 | 3 | 3 | 1 | 211 | 23,757 | 6,7 | Not Found | Not Found | High | High | 0,002 | 96,46 | 3 |
| O14654 | IRS4 | Insulin receptor substrate 4 OS=Homo sapiens GN=IRS4 PE=1 SV=1 | 1,272871917 | 1 | 1 | 1 | 1 | 1257 | 133,685 | 8,44 | Not Found | Not Found | High | High | 0,003 | 93,65 | 1 |
| Q9BQ70 | TCF25 | Transcription factor 25 OS=Homo sapiens GN=TCF25 PE=1 SV=1 | 6,952662722 | 2 | 2 | 2 | 1 | 676 | 76,619 | 6,35 | Not Found | Not Found | High | High | 0,003 | 93,08 | 2 |
| Q13751 | LAMB3 | Laminin subunit beta-3 OS=Homo sapiens GN=LAMB3 PE=1 SV=1 | 1,279863481 | 1 | 1 | 1 | 1 | 1172 | 129,489 | 7,21 | Not Found | Not Found | High | High | 0,003 | 92,70 | 1 |
| Q13263 | TRIM28 | Transcription intermediary factor 1-beta OS=Homo sapiens GN=TRIM28 PE=1 SV=5 | 5,508982036 | 2 | 2 | 2 | 1 | 835 | 88,493 | 5,77 | Not Found | Not Found | High | High | 0,003 | 90,57 | 2 |
| Q7Z460 | CLASP1 | CLIP-associating protein 1 OS=Homo sapiens GN=CLASP1 PE=1 SV=1 | 1,88556567 | 2 | 2 | 2 | 1 | 1538 | 169,346 | 9,03 | Not Found | Not Found | High | High | 0,004 | 85,68 | 2 |
| Q8NDV7 | TNRC6A | Trinucleotide repeat-containing gene 6A protein OS=Homo sapiens GN=TNRC6A PE=1 SV=2 | 2,650356779 | 4 | 4 | 4 | 1 | 1962 | 210,169 | 7,01 | Not Found | Not Found | High | High | 0,004 | 83,93 | 4 |
| P42357 | HAL | Histidine ammonia-lyase OS=Homo sapiens GN=HAL PE=1 SV=1 | 6,849315068 | 2 | 2 | 2 | 1 | 657 | 72,652 | 6,95 | Not Found | Not Found | High | High | 0,004 | 79,82 | 2 |
| Q99569 | PKP4 | Plakophilin-4 OS=Homo sapiens GN=PKP4 PE=1 SV=2 | 1,929530201 | 2 | 2 | 2 | 1 | 1192 | 131,787 | 8,94 | Not Found | Not Found | High | High | 0,004 | 78,86 | 2 |
| A7E2V4 | ZSWIM8 | Zinc finger SWIM domain-containing protein 8 OS=Homo sapiens GN=ZSWIM8 PE=1 SV=1 | 1,741970604 | 2 | 2 | 2 | 1 | 1837 | 197,173 | 6,8 | Not Found | Not Found | High | High | 0,004 | 75,84 | 2 |
| Q8IWT3 | CUL9 | Cullin-9 OS=Homo sapiens GN=CUL9 PE=1 SV=2 | 1,470003973 | 3 | 3 | 3 | 1 | 2517 | 281,049 | 5,45 | Not Found | Not Found | Medium | High | 0,004 | 75,06 | 3 |
| Q5VUA4 | ZNF318 | Zinc finger protein 318 OS=Homo sapiens GN=ZNF318 PE=1 SV=2 | 1,360245722 | 2 | 2 | 2 | 1 | 2279 | 250,958 | 7,2 | Not Found | Not Found | High | High | 0,004 | 73,88 | 2 |
| P38432 | COIL | Coilin OS=Homo sapiens GN=COIL PE=1 SV=1 | 1,736111111 | 1 | 2 | 1 | 1 | 576 | 62,57 | 9,07 | Not Found | Not Found | High | High | 0,004 | 73,03 | 1 |
| Q8WV41 | SNX33 | Sorting nexin-33 OS=Homo sapiens GN=SNX33 PE=1 SV=1 | 5,226480836 | 2 | 2 | 2 | 1 | 574 | 65,223 | 6,79 | Not Found | Not Found | High | High | 0,004 | 67,40 | 2 |
| Q9BQS8 | FYCO1 | FYVE and coiled-coil domain-containing protein 1 OS=Homo sapiens GN=FYCO1 PE=1 SV=3 | 0,947225981 | 1 | 1 | 1 | 1 | 1478 | 166,879 | 4,92 | Not Found | Not Found | High | High | 0,004 | 65,30 | 1 |
| Q641Q2 | WASHC2A | WASH complex subunit 2A OS=Homo sapiens GN=WASHC2A PE=1 SV=3 | 1,640566741 | 1 | 1 | 1 | 1 | 1341 | 147,095 | 4,81 | Not Found | Not Found | High | High | 0,010 | 60,50 | 1 |
| Q15154 | PCM1 | Pericentriolar material 1 protein OS=Homo sapiens GN=PCM1 PE=1 SV=4 | 1,333992095 | 2 | 2 | 2 | 1 | 2024 | 228,392 | 5,02 | Not Found | Not Found | Medium | High | 0,010 | 60,23 | 2 |
| Q9Y5Q9 | GTF3C3 | General transcription factor 3C polypeptide 3 OS=Homo sapiens GN=GTF3C3 PE=1 SV=1 | 1,58013544 | 1 | 1 | 1 | 1 | 886 | 101,208 | 5,07 | Not Found | Not Found | High | Medium | 0,010 | 56,61 | 1 |

| | | | | | | | | | | | | | | | | | |
|---|---|---|---|---|---|---|---|---|---|---|---|---|---|---|---|---|---|
| Q9UPW5 | AGTPBP1 | Cytosolic carboxypeptidase 1 OS=Homo sapiens GN=AGTPBP1 PE=1 SV=3 | 0,897226754 | 1 | 1 | 1 | 1 | 1226 | 138,361 | 6,15 | Not Found | Not Found | High | Medium | 0,011 | 55,71 | 1 |
| Q9HCJ0 | TNRC6C | Trinucleotide repeat-containing gene 6C protein OS=Homo sapiens GN=TNRC6C PE=1 SV=3 | 0,828402367 | 1 | 1 | 1 | 1 | 1690 | 175,857 | 6,96 | Not Found | Not Found | High | Medium | 0,012 | 54,00 | 1 |
| Q16787 | LAMA3 | Laminin subunit alpha-3 OS=Homo sapiens GN=LAMA3 PE=1 SV=2 | 0,420042004 | 1 | 1 | 1 | 1 | 3333 | 366,414 | 7,24 | Not Found | Not Found | High | Medium | 0,013 | 53,17 | 1 |
| O00468 | AGRN | Agrin OS=Homo sapiens GN=AGRN PE=1 SV=5 | 0,53217223 | 1 | 1 | 1 | 1 | 2067 | 217,092 | 6,39 | Not Found | Not Found | High | Medium | 0,013 | 52,88 | 1 |
| P07942 | LAMB1 | Laminin subunit beta-1 OS=Homo sapiens GN=LAMB1 PE=1 SV=2 | 1,231802912 | 1 | 1 | 1 | 1 | 1786 | 197,909 | 4,94 | Not Found | Not Found | High | Medium | 0,013 | 52,29 | 1 |
| P31944 | CASP14 | Caspase-14 OS=Homo sapiens GN=CASP14 PE=1 SV=2 | 5,785123967 | 1 | 1 | 1 | 1 | 242 | 27,662 | 5,58 | Not Found | Not Found | High | Medium | 0,013 | 51,11 | 1 |
| P62847 | RPS24 | 40S ribosomal protein S24 OS=Homo sapiens GN=RPS24 PE=1 SV=1 | 9,022556391 | 1 | 1 | 1 | 1 | 133 | 15,413 | 10,78 | Not Found | Not Found | High | Medium | 0,013 | 50,81 | 1 |
| P36578 | RPL4 | 60S ribosomal protein L4 OS=Homo sapiens GN=RPL4 PE=1 SV=5 | 2,576112412 | 1 | 1 | 1 | 1 | 427 | 47,667 | 11,06 | Not Found | Not Found | High | Medium | 0,013 | 50,62 | 1 |
| Q6UWP8 | SBSN | Suprabasin OS=Homo sapiens GN=SBSN PE=1 SV=2 | 3,050847458 | 1 | 1 | 1 | 1 | 590 | 60,505 | 7,01 | Not Found | Not Found | High | Medium | 0,013 | 50,43 | 1 |
| P19823 | ITIH2 | Inter-alpha-trypsin inhibitor heavy chain H2 OS=Homo sapiens GN=ITIH2 PE=1 SV=2 | 1,585623679 | 1 | 1 | 1 | 1 | 946 | 106,397 | 6,86 | Not Found | Not Found | High | Medium | 0,013 | 47,99 | 1 |
| P04004 | VTN | Vitronectin OS=Homo sapiens GN=VTN PE=1 SV=1 | 3,138075314 | 1 | 1 | 1 | 1 | 478 | 54,271 | 5,8 | Not Found | Not Found | High | Medium | 0,013 | 46,58 | 1 |
| O43795 | MYO1B | Unconventional myosin-Ib OS=Homo sapiens GN=MYO1B PE=1 SV=3 | 1,848591549 | 1 | 1 | 1 | 1 | 1136 | 131,902 | 9,38 | Not Found | Not Found | Medium | Medium | 0,016 | 43,67 | 1 |
| P21333 | FLNA | Filamin-A OS=Homo sapiens GN=FLNA PE=1 SV=4 | 0,491122025 | 1 | 1 | 1 | 1 | 2647 | 280,564 | 6,06 | Not Found | Not Found | Medium | Medium | 0,016 | 43,12 | 1 |
| Q9BWT7 | CARD10 | Caspase recruitment domain-containing protein 10 OS=Homo sapiens GN=CARD10 PE=1 SV=2 | 1,937984496 | 1 | 1 | 1 | 1 | 1032 | 115,859 | 5,95 | Not Found | Not Found | Medium | Medium | 0,017 | 42,85 | 1 |
| A2RUB1 | MEIOC | Meiosis-specific coiled-coil domain-containing protein MEIOC OS=Homo sapiens GN=MEIOC PE=2 SV=3 | 1,470588235 | 1 | 1 | 1 | 1 | 952 | 107,491 | 7,12 | Not Found | Not Found | Medium | Medium | 0,021 | 39,82 | 1 |
| Q02978 | SLC25A11 | Mitochondrial 2-oxoglutarate/malate carrier protein OS=Homo sapiens GN=SLC25A11 PE=1 SV=3 | 5,095541401 | 1 | 1 | 1 | 1 | 314 | 34,04 | 9,91 | Not Found | Not Found | Medium | Medium | 0,023 | 38,81 | 1 |
| Q5JTC6 | AMER1 | APC membrane recruitment protein 1 OS=Homo sapiens GN=AMER1 PE=1 SV=2 | 1,674008811 | 1 | 1 | 1 | 1 | 1135 | 123,952 | 4,84 | Not Found | Not Found | Medium | Medium | 0,023 | 38,28 | 1 |
| Q9UDY2 | TJP2 | Tight junction protein ZO-2 OS=Homo sapiens GN=TJP2 PE=1 SV=2 | 1,428571429 | 1 | 1 | 1 | 1 | 1190 | 133,876 | 7,4 | Not Found | Not Found | Medium | Medium | 0,027 | 37,35 | 1 |
| Q9UHR4 | BAIAP2L1 | Brain-specific angiogenesis inhibitor 1-associated protein 2-like protein 1 OS=Homo sapiens GN=BAIAP2L1 PE=1 SV=2 | 2,935420744 | 1 | 1 | 1 | 1 | 511 | 56,847 | 8,68 | Not Found | Not Found | Medium | Medium | 0,030 | 35,57 | 1 |
| P17040 | ZSCAN20 | Zinc finger and SCAN domain-containing protein 20 OS=Homo sapiens GN=ZSCAN20 PE=2 SV=3 | 1,821668265 | 1 | 1 | 1 | 1 | 1043 | 117,467 | 6,43 | Not Found | Not Found | Medium | Medium | 0,041 | 34,29 | 1 |
| Q5T0W9 | FAM83B | Protein FAM83B OS=Homo sapiens GN=FAM83B PE=1 SV=1 | 2,176063304 | 1 | 1 | 1 | 1 | 1011 | 114,729 | 8,97 | Not Found | Not Found | Medium | Medium | 0,047 | 34,04 | 1 |
| A3KN83 | SBNO1 | Protein strawberry notch homolog 1 OS=Homo sapiens GN=SBNO1 PE=1 SV=1 | 0,502512563 | 1 | 1 | 1 | 1 | 1393 | 154,216 | 7,88 | Not Found | Not Found | Medium | Medium | 0,048 | 33,99 | 1 |
| Q5TGY3 | AHDC1 | AT-hook DNA-binding motif-containing protein 1 OS=Homo sapiens GN=AHDC1 PE=1 SV=1 | 0,748596382 | 1 | 1 | 1 | 1 | 1603 | 168,245 | 9,04 | Not Found | Not Found | Medium | Medium | 0,048 | 33,70 | 1 |
| P42766 | RPL35 | 60S ribosomal protein L35 OS=Homo sapiens GN=RPL35 PE=1 SV=2 | 8,130081301 | 1 | 1 | 1 | 1 | 123 | 14,543 | 11,05 | Not Found | Not Found | Medium | Medium | 0,048 | 33,59 | 1 |
| P84090 | ERH | Enhancer of rudimentary homolog OS=Homo sapiens GN=ERH PE=1 SV=1 | 10,57692308 | 1 | 1 | 1 | 1 | 104 | 12,251 | 5,92 | Not Found | Not Found | Medium | Medium | 0,048 | 33,47 | 1 |

**For both Supplementary Table 1 and 2:**
Shown are
-proteins that were detected as "High" or "Medium"in the "E1B-55k Ab" sampe and "not found "in the "Beads only"  nor "Mouse IgG"control samples.
-proteins with "Protein FDR Confidence Mascot" "High" or "Medium".

The proteins are sorted based on their "Mascot Score".
